# Clathrin mediates both internalization and vesicular release of triggered T cell receptor at the immunological synapse through distinct adaptors

**DOI:** 10.1101/2022.02.02.478780

**Authors:** Audun Kvalvaag, Pablo F Céspedes, Salvatore Valvo, David G Saliba, Elke Kurz, Kseniya Korobchevskaya, Michael L Dustin

## Abstract

Ligation of the T cell receptor (TCR) to peptide-MHC complexes initiates signaling leading to T cell activation. Regulation of T cell responses also requires mechanisms to stop this signaling and to downregulate surface expression of the receptor. T cells achieve this both by TCR internalization and by releasing TCR loaded vesicles directly from the plasma membrane. How these distinct fates are coordinated is unknown. Here we show that clathrin is recruited to TCR microclusters by HRS and STAM2 and that this process is essential for TCR release. Subsequently, EPN1 recruits clathrin to the remaining antigen conjugated TCRs to enable a late wave of TCR endocytosis. With these results we demonstrate two clathrin-dependent mechanisms and show how the clathrin machinery participates in membrane evagination and invagination depending upon the adaptor recruiting it. These sequential mechanisms mediate bi-directional membrane exchange at the immunological synapse, providing a scaffold for critical communication.

## Introduction

The fundamental molecular interactions responsible for regulating the adaptive immune response occur within a nanoscale gap between T cells and antigen presenting cells (APCs), termed the immunological synapse (IS). IS formation is induced upon TCR interactions with agonist peptide-MHC (pMHC) on the surface of APCs ^1^. This process can be recapitulated by antigen presentation on supported lipid bilayers (SLBs), a minimal component system composed of a mobile lipid phase configured to carry proteins at any given density ^2^.

During activation, the components of the synapse rearrange in the opposing lipid membranes to form a characteristic bullseye pattern consisting of three primary domains ^3,4^. The bullseye itself is termed the central supramolecular activation cluster (cSMAC) and is dominated by TCR and associated pMHC. This area is surrounded by the peripheral supramolecular activation cluster (pSMAC) which forms an adhesive ring defined by LFA-1 on the T cell side associated with ICAM-1 on the bilayer side. Beyond the pSMAC is the distal supramolecular activation cluster (dSMAC) which is dominated by filamentous actin (F-actin) and an adhesion corolla ^5^. TCR engagement is initiated in microclusters that arise from filopodia in the dSMAC and retain protrusive activity even as they traffic through the pSMAC toward the cSMAC ^6–9^.

The cSMAC was originally thought to be a specialized domain for TCR signal enhancement and receptor downregulation through internalization of activated TCRs ^10^. However, Choudhuri *et al*. discovered in 2014 that TCRs can also be incorporated in vesicles released directly from the plasma membrane into the extracellular space. Whereas extracellular vesicles released by exocytosis of multivesicular bodies are called exosomes, vesicles that are released directly from the plasma membrane are termed ectosomes ^11^. Therefore, we have defined vesicles released directly into the immune synaptic cleft as synaptic ectosomes ^12^. These are formed in a process requiring the ESCRT-I component TSG101 and the ESCRT-III associated ATPase Vps4. This is analogous to the process occurring on multivesicular bodies during formation of intraluminal vesicles (ILVs) and during HIV budding from the plasma membrane ^13,14^. During ILV formation, the ESCRT-0 components HRS and STAM2 recruit clathrin, which in turn assembles into flat lattices to mediate recruitment of subsequent ESCRT machinery and ubiquitinated receptors ^15,16^. However, it is not known if clathrin is involved in the formation of synaptic ectosomes.

Clathrin is primarily known for its role in endocytosis, where it is recruited to the plasma membrane by adaptor proteins such as EPN1 (also called epsin-1) and AP2. Together, these proteins facilitate deformation and invagination of the membrane which is ultimately pinched off as a clathrin coated endocytic vesicle by the large GTPase dynamin ^17^. Previous reports have shown that this mechanism is engaged to internalize TCR during constitutive TCR recycling ^18,19^ and to internalize non-activated bystander TCR during T cell activation, while it has been suggested that clathrin is not involved in downregulating triggered TCR ^20–22^. Clathrin and HRS positive vesicles have also been shown to polarize towards the immunological synapse during T cell activation, and they have been implicated in recruiting actin there ^23^. However, these studies were performed under a paradigm in which the ability of T cells to release TCR from the plasma membrane in synaptic ectosomes was not recognized.

Here, we show that clathrin is pivotal during T cell activation through its essential role in two sequential processes. First, clathrin is essential for ESCRT-mediated release of TCR loaded synaptic ectosomes at the cSMAC. As the synapse matures, there is a temporal switch from the clathrin-associated ESCRT components HRS and STAM2 to the endocytic clathrin adaptor EPN1. Remaining antigen-ligated TCR are then internalized via clathrin coated vesicles.

## Results

### Clathrin is recruited to TCR microclusters at the immunological synapse

The immunological synapse undergoes several stages of maturation during T cell activation on SLBs, as illustrated in Figure 1a (inserts depict TCR microcluster formation and movement at the SLB as seen by an inverted microscope). The process is initiated during presynaptic interactions of TCR at the tips of microvilli with antigen leading to rapid activation of LFA-1-ICAM-1 mediated adhesion and immediately followed by spreading of the T cell with the formation of LFA-1 and TCR microclusters (0-5 min). LFA-1 and TCR then move centripetally in the nascent synapse and accumulate in the pSMAC and cSMAC, respectively, of the maturing synapse (5-20 min). After about 30 minutes signaling is sustained by a minimal number of TCR microclusters and formation of a motile kinapse is often observed (>30 min) ^24,25^.

**Figure 1.**
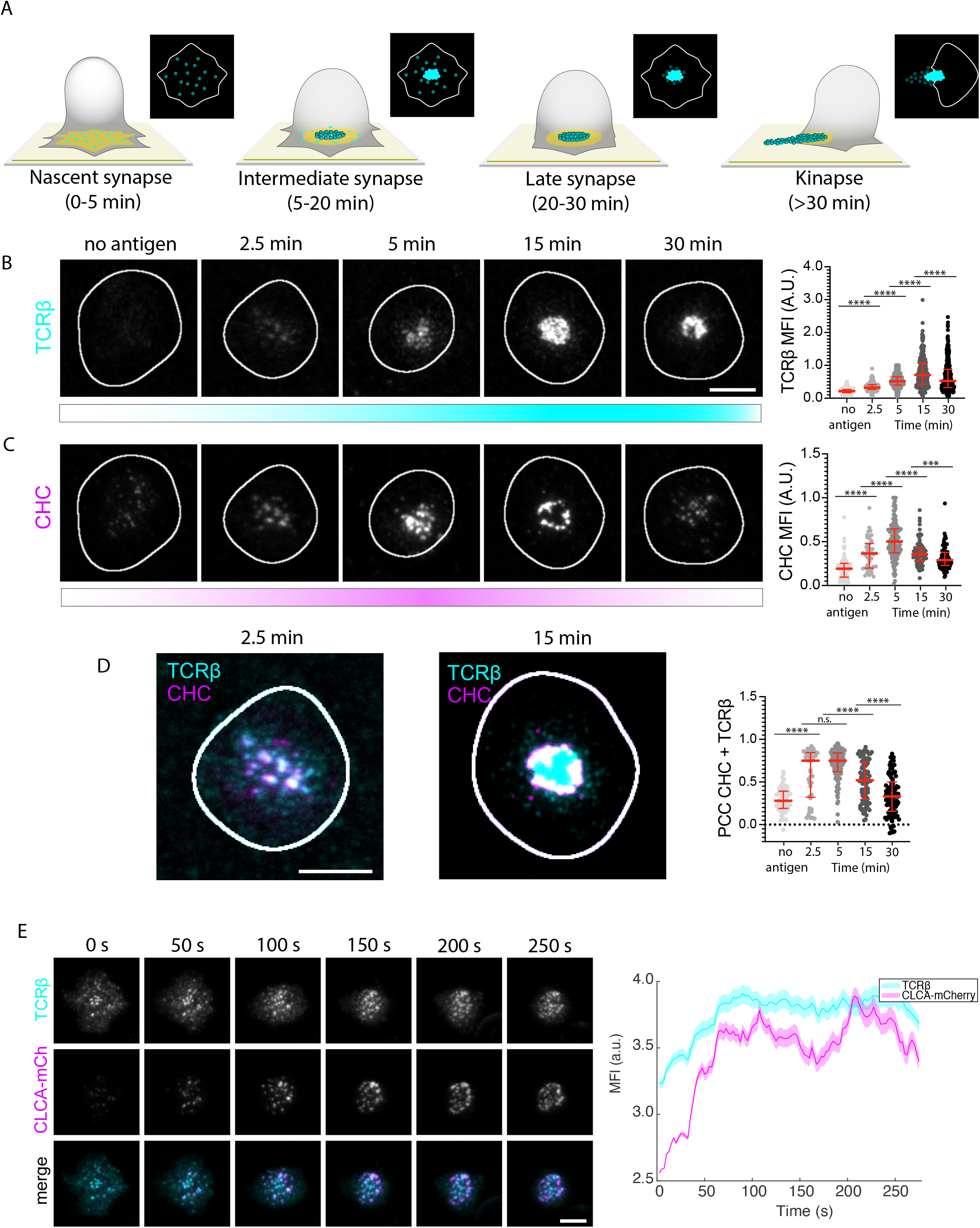
Clathrin is recruited to the immunological synapse. (**A**) Schematic of the different maturation stages of the immunological synapse formed between a T cell and an SLB. ICAM-1 (yellow) forms the adhesion ring and TCRβ (cyan) microclusters reach the center of the contact area, where some are released as synaptic ectosomes that are left behind as symmetry breaking allows the T cell to relocate. (**B-C**) Representative TIRF micrographs of AND mCD4 T cells incubated on SLBs either with ICAM-1-AF405 (200/μm^2^) alone for 5 min or with ICAM-1-AF405 + I-E^k^-MCC (20/μm^2^) for 2.5, 5, 15 and 30 min, labelled with anti-mouse TCRβ (cyan) and anti-CHC (magenta). N_cells_ ≥ 45 per timepoint. Scale bar, 5 μm. The right panels are quantifications of the mean fluorescence intensity (MFI) of TCRβ and CHC. ***, p value < 0.001; ****, p value of < 0.0001 with unpaired two-tailed non-parametric Mann Whitney test. Lines represent median value and error bars represent interquartile range. (**D**) Representative micrographs emphasizing the colocalization between CHC (magenta) and TCRβ (cyan) at 2.5 min and 15 min. The right panel is quantification of the Pearson Correlation Coefficient (PCC) between CHC and TCRβ across the synaptic interphase from the micrographs in (**B-C**). ****, p value of < 0.0001 with unpaired two-tailed non-parametric Mann Whitney test. Lines represent median value and error bars represent interquartile range. (**E**) Representative timeframes from a movie of a live mCD4 AND T cell expressing CLCA-mCherry (magenta) while forming an immunological synapse on an SLB with ICAM-1-AF405 (200/μm^2^) and I-E^k^-MCC (50/μm^2^). The TCR is labelled with anti-TCRβ (cyan). The right panel is mean temporal fluorescence intensity traces +/− SEM of TCR microclusters with overlapping CLCA-mCherry fluorescence. N_cells_ = 5.

We first asked if clathrin is recruited to TCR microclusters at any stage during immunological synapse formation. Monoclonal T cells from AND TCR transgenic mice specific for I-E^k^ with a moth cytochrome C peptide 88-103 (I-E^k^-MCC) were incubated on SLB with ICAM-1-AF405 (200 μm^−2^) alone for 5 minutes, or with ICAM-1 plus I-E^k^-MCC (20 μm^-2^) for 2.5, 5, 15 and 30 minutes. We then fixed the T cells and immunolabelled the TCR with the anti-mouse TCRβ Fab H57 tagged with Alexa Flour 488 (TCRβ) and clathrin with an anti-clathrin heavy chain (CHC) antibody. By imaging the cells by total internal reflection fluorescence (TIRF) microscopy, we observed formation of TCR microclusters after 2.5 minutes in the presence of antigen and an increase in TCRβ fluorescence intensity compared to no antigen (Figure 1B). The TCRβ intensity peaked after 15 minutes before decreasing by about 50% after 30 minutes. Intriguingly, the increase in TCRβ intensity after 2.5 minutes was accompanied by a two-fold increase in CHC fluorescence intensity (Figure 1C). This correlated with increased colocalization between CHC and TCR microclusters, with a median Pearson Correlation Coefficient (PCC) of 0.75 at 2.5 minutes and 0.73 at 5 minutes. The CHC intensity peaked after 5 minutes and declined as the synapse matured and TCR accumulated in the cSMAC. After 15 minutes, clathrin appeared to be confined to the cSMAC periphery and at 30 minutes clathrin was no longer strongly associated with TCR in the synapse (Figure 1D). By increasing the I-E^k^-MCC density on the bilayer, we could also observe that the CHC intensity increased accordingly (Figure S1A). Live imaging of synapses with mCherry-tagged clathrin light chain A (CLCA-mCherry) ^26^ revealed that TCR and CLCA recruitment increased as microclusters moved centripetally (Figure 1E, Movie S1). CHC recruitment to TCR microclusters was also observed in human CD4 (hCD4) T cells on SLBs with ICAM-1 and an agonistic anti-CD3ε Fab UCHT1 with a PCC of 0.82 after 5 minutes (Figure S1B). We conclude that clathrin is recruited to TCR microclusters as they move to the center of the nascent synapse, but that it is largely excluded from the center of the mature cSMAC.

### Clathrin adaptors recruited to the synapse undergo a temporal shift from the ESCRT-0 components HRS and STAM2, to the endocytic adaptor EPN1

Previous reports have shown that the ESCRT-I component TSG101 is required for cSMAC formation whereas the ESCRT associated ATPase VPS4 is required for scission of the bud-neck of TCR loaded nascent synaptic ectosomes ^12,27^. This process is analogous to ESCRT-mediated formation of ILVs in which TSG101 is recruited to flat clathrin lattices on the limiting endosomal membrane by HRS and STAM2 ^16^. We therefore asked whether a similar mechanism was taking place at the immunological synapse.

We incubated T cells on bilayers with ICAM-1 alone for 5 min or with ICAM-1 plus I-E^k^-MCC for 2.5, 5, 15 and 30 minutes before we fixed and permeabilized them. We then immunolabelled HRS and STAM2 and could observe potent recruitment of both proteins in response to antigen already after 2.5 minutes with a twofold increase in MFI compared to no antigen (Figure 2A-B and S2A-B). As for clathrin, HRS and STAM2 recruitment peaked after 5 minutes before the fluorescence intensity dropped after 15 minutes. The early recruitment correlated with extensive colocalization with TCR microclusters with median PCCs of 0.79 and 0.77 (HRS) and 0.74 and 0.69 (STAM2) at 2.5 and 5 minutes, respectively. This was reduced to 0.49 and 0.44 at 15 minutes and 0.45 and 0.42 at 30 minutes (Figure 2C and S2C).

**Figure 2.**
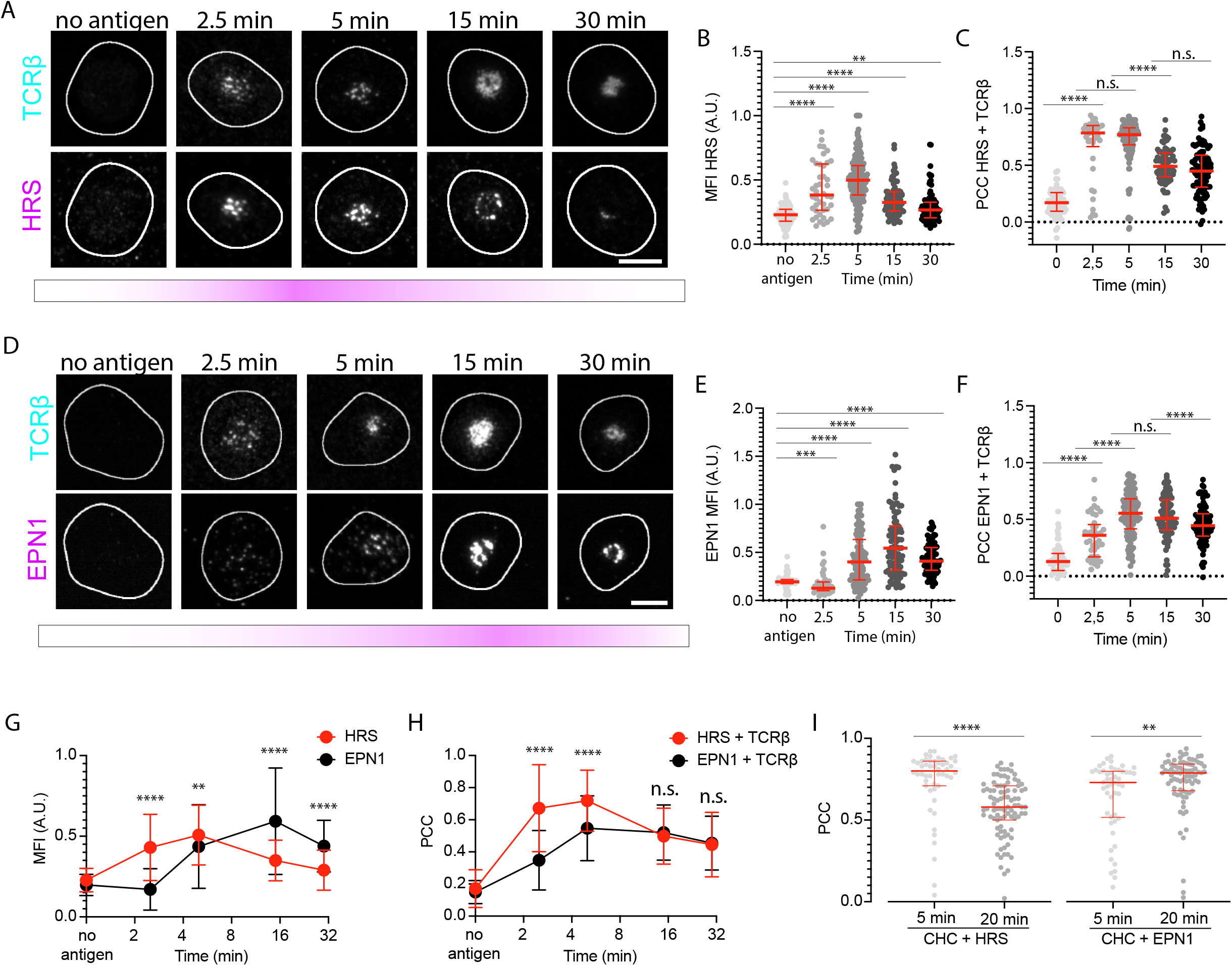
HRS and EPN1 are recruited to the immunological synapse at different stages of synapse maturation. (**A**) Representative TIRF micrographs of AND mCD4 T cells incubated on SLBs either with ICAM-1-AF405 (200/μm^2^) alone for 5 min or with ICAM-1-AF405 + I-E^k^-MCC (20/μm^2^) for 2.5, 5, 15 and 30 min, labelled with anti-mouse TCRβ (cyan) and anti-HRS (magenta). N_cells_ ≥ 42 per timepoint. Scale bar, 5 μm. (**B-C**) Quantification of the temporal MFI of HRS (**B**) and the temporal PCC between HRS and TCRβ (**C**) across the synaptic interphase from the micrographs in (**A**). **, p value < 0.01; ****, p value of < 0.0001 with unpaired two-tailed non-parametric Mann Whitney test. Lines represent median value and error bars represent interquartile range (**D**) Representative TIRF micrographs of AND mCD4 T cells incubated on SLBs either with ICAM-1-AF405 (200/μm^2^) alone for 5 min or with ICAM-1-AF405 + I-E^k^-MCC (20/μm^2^) for 2.5, 5, 15 and 30 min, labelled with anti-mouse TCRβ (cyan) and anti-EPN1 (magenta). N_cells_ ≥ 45 per timepoint. Scale bar, 5 μm. (**E-F**) Quantification of the temporal MFI of EPN1 (**E**) and the temporal PCC between EPN1 and TCRβ (**F**) across the synaptic interphase from the micrographs in (**D**). ***, p value < 0.001; ****, p value of < 0.0001 with unpaired two-tailed non-parametric Mann Whitney test. Lines represent median value and error bars represent interquartile range. (**G-H**) Direct comparison of the temporal MFI +/− SD of HRS and EPN1 at 2.5, 5, 15 and 30 min relative to the max intensity of each protein at 5 min (**G**) and the temporal PCC between HRS and TCRβ compared to the temporal PCC of EPN1 and TCRβ (**H**) from the micrographs in (**A**) and (**D**). **, p value < 0.01; ****, p value of < 0.0001 with unpaired two-tailed non-parametric Mann Whitney test. (**I**) Temporal PCC between HRS and CHC, and EPN1 and CHC. N_cells_ ≥ 49 per timepoint. **, p value < 0.01; ****, p value of < 0.0001 with unpaired two-tailed non-parametric Mann Whitney test. Lines represent median value and error bars represent interquartile range.

We have also previously reported that the endocytic clathrin adaptor EPN1 localizes to the immunological synapse during T cell activation ^28^. Here, we confirm those observations and show that as for HRS, STAM2 and CHC, this recruitment depends on TCR-antigen ligation (Figure 2D). However, while HRS and STAM2 were recruited already after 2.5 minutes and peaked after 5 minutes, we could detect a slight decrease in EPN1 MFI at 2.5 minutes, before it increased at 5 minutes. EPN1 recruitment then peaked after 15 minutes with a threefold increase in MFI compared to no antigen (Figure 2E). EPN1 colocalized with TCR with a PCC of 0.36 at 2.5 minutes, 0.6 at 5 minutes, 0.57 at 15 minutes and 0.43 at 30 minutes (Figure 2F). When directly compared to HRS recruitment, we could observe significantly less EPN1 recruitment at 2.5 minutes and 5 minutes relative to maximum MFI at 5 minutes compared to HRS. However, after 15 and 30 minutes this was reversed with significantly more EPN1 recruited than HRS (Figure 2G). In terms of colocalization with TCR, this was significantly higher for HRS after 2.5 and 5 minutes, but at the same level for HRS and EPN1 after 15 and 30 minutes (Figure 2H).

When we immunostained HRS, STAM2 and EPN1 together with CHC, we could observe a high degree of colocalization between these proteins and clathrin at 5 minutes with PCCs of 0.8, 0.77 and 0.75, respectively (Figure 2I and S2D). However, for HRS and STAM2 this was reduced to 0.55 and 0.57 after 20 minutes, while it increased to 0.8 for EPN1.

AP2 is regarded as the primary endocytic clathrin adaptor and is an integral part of the majority of clathrin coated pits ^29^. We therefore analyzed temporal AP2 recruitment to the immunological synapse as well and could observe a significant increase in response to antigen stimulation (Figure S2E-F). However, this increase did not correlate with increased colocalization with TCR microclusters which was essentially non-existent across all timepoints (Figure S2G).

Taken together, these results show that clathrin adaptor recruitment to the immunological synapse is regulated in a temporal manner, initially being dominated by the ESCRT-0 components HRS and STAM2, and subsequently the endocytic adaptor protein EPN1. AP2 is recruited to the plasma membrane in response to antigen, but not to TCR microclusters.

### TSG101, HRS and clathrin are required for cSMAC formation

We next depleted CHC, HRS, EPN1, AP2 and STAM2 by CRISPR/Cas9 mediated knockout (KO) in hCD4 T cells and analyzed immunological synapse formation on bilayers with UCHT1-AF488 (30 μm^−2^) and ICAM1-AF405 (200 μm^−2^) after 5 and 20 minutes. CD19 and TSG101 KO were included as negative and positive controls, respectively (Figure S3A). To analyze the synaptic distribution of ICAM1, CHC and UCHT1 in response to the knockouts, we segmented the cells and performed radial averaging on them (Figure 3A-B). We could then observe defective cSMAC formation following clathrin, HRS, STAM2 and TSG101 KO, with UCHT1 labeled CD3ε appearing unable to fully translocate to the center of the contact area and ICAM-1 being incompletely excluded from the cSMAC after 20 min as reported following siRNA-mediated TSG101 knockdown in T cells previously ^27^ (Figure 3C, Figure S3B). For HRS and TSG101 KO, this was accompanied by a strong increase in cSMAC localized CHC, while for CHC KO, the ICAM-1 ring seemed to collapse into the cSMAC after 20 minutes incubation on SLBs.

**Figure 3.**
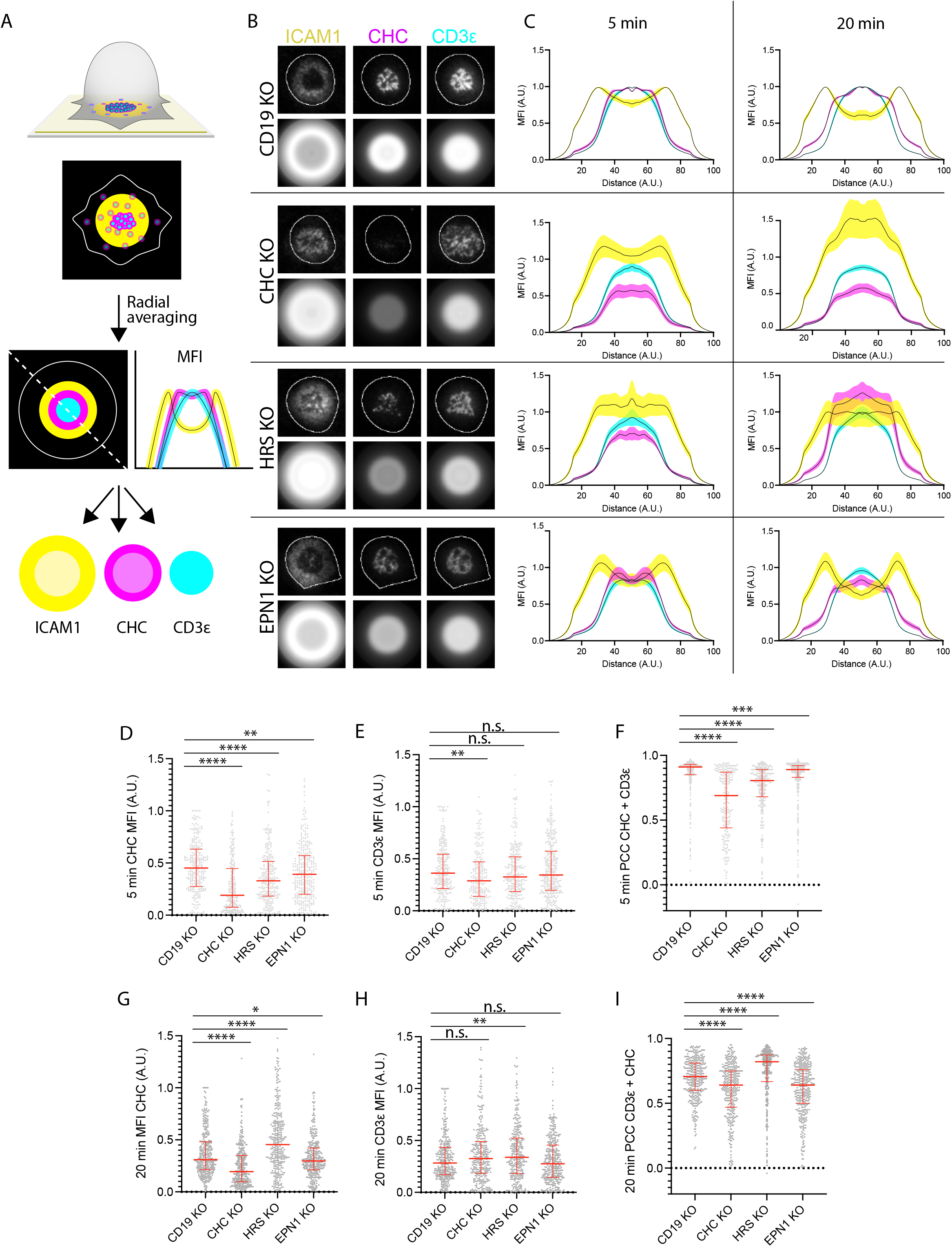
HRS, STAM2 and EPN1 sequentially recruit clathrin to the immunological synapse. (**A**) Schematic of radial averaging of micrographs of the immunological synapse formed between a T cell and a SLB. ICAM-1 is labelled yellow, TCR cyan and CHC magenta. The dashed line and the line plots represent a diagonal measurement of the positional mean fluorescence intensity of the individual channels. (**B**) Representative TIRF micrographs and corresponding radial averages of hCD4 CD19, CHC, HRS and EPN1 KO T cells incubated on SLBs with ICAM-1-AF405 (200/μm^2^) and UCHT1-AF488 (CD3ε, 30/μm^2^) for 5 min. (**C**) Radial averages of hCD4 CD19, CHC, HRS and EPN1 KO T cells incubated on the bilayers from (**B**) for 5 and 20 minutes. The MFI represents mean fluorescence intensity from 5 individual experiments +/− SEM. (**D-I**) Quantification of the MFI of CHC (**D, G**), CD3ε (**E, H**) and the PCC between CHC and CD3ε (**F, I**) across the synaptic interphase from hCD4 CD19, CHC, HRS and EPN1 KO T cells incubated on the bilayers from (**B**) for 5 and 20 minutes, respectively. *, p-value of < 0.05; **, p-value of < 0.01; ***, p value of < 0.001; ****, p value of < 0.0001 with unpaired twotailed non-parametric Mann Whitney test. N_cells_ ≥ 219 per condition. Lines represent median value and error bars represent interquartile range.

When we then examined clathrin recruitment to the synaptic interphase in these cells, we observed a significant drop following HRS, EPN1 and STAM2 KO after 5 minutes incubation on bilayers, thus confirming that these proteins are indeed recruiting clathrin there at this timepoint (Figure 3D, Figure S3C). We could also detect a drop in CD3ε MFI after 5 minutes following CHC KO, which might indicate reduced TCR expression at the plasma membrane following CHC KO, possibly due to defective TCR recycling (Figure 3E). When we analyzed colocalization between CHC and CD3ε after 5 minutes, we could observe a strong decrease following HRS KO compared to the other conditions (Figure 3F, Figure S3D). While there was also a significant decrease following EPN1, STAM2 and TSG101 KO, these data suggest that HRS is the primary adaptor protein for recruiting clathrin to TCR microclusters early during synapse formation.

Intriguingly, clathrin recruitment was strongly increased after 20 minutes following HRS and TSG101 KO, while we observed a slight decrease following EPN1 and STAM2 KO (Figure 3G, Figure S3E). We observed an increase in CD3ε MFI following AP2, HRS and TSG101 KO as reported previously following siRNA knockdown of HRS and TSG101 in AND T cells ^27^ (Figure 3H, Figure S3F). When we examined colocalization between CHC and UCHT1, we observed a strong decrease following EPN1 KO, while there was a strong increase following HRS KO and TSG101 KO (Figure 3I, Figure S3G).

These data suggest that EPN1 is the primary adaptor protein for recruiting clathrin to the mature cSMAC, while HRS and TSG101 KO seem to induce defective clearance of activated TCR from the membrane and clathrin arrest. Interestingly, STAM2 KO seems to decrease clathrin recruitment both early and late during synapse formation, thus indicating that it not only acts as a partner for HRS in the ESCRT-0 complex in this context, but also has an additional role during late-stage immunological synapse formation. AP2 KO did not affect clathrin recruitment at any time, but the slight increase in CD3ε MFI after 20 minutes might imply that TCR/CD3 surface expression had increased due to diminished steady state turnover, as this has been reported to be mediated by AP2 and clathrin previously ^30^.

We next investigated clathrin and CD3ε in the mature cSMAC following CD19, CHC and HRS KO by extended Total Internal Reflection Fluorescence Structured Illumination Microscopy (eTIRF-SIM) with up to 90 nm axial and 150 nm lateral resolution (Figure S3H). We could then clearly observe CHC at the edge of the cSMAC after 20 minutes. However, while for the CD19 KO cells the CD3ε pattern appeared highly punctuate, it appeared as an interconnected web-like structure in the CHC and HRS KO cells. The punctuate pattern observed following CD19 KO might thus represent released TCR loaded synaptic ectosomes, while the web-like pattern observed following CHC and HRS KO might represent CD3/TCR arrested in the plasma membrane.

### HRS and clathrin are required for transfer of TCR loaded vesicles

To examine whether these proteins are indeed involved in release of TCR loaded vesicles, we next investigated TCR transfer to bead supported lipid bilayers (BSLBs) as illustrated in Figure 4A. This was done by incubating human CD4 T cells with bilayer coated silica beads at a 1:1 ratio for 90 minutes before detaching them and analyzing them by flow cytometry (described in Saliba *et al*. ^28^). We could then observe a 50% reduction in the amount of TCR transferred to the beads following CHC and HRS KO (Figure 4B). These data demonstrate that clathrin and HRS are essential for transfer of TCR at the immunological synapse.

**Figure 4.**
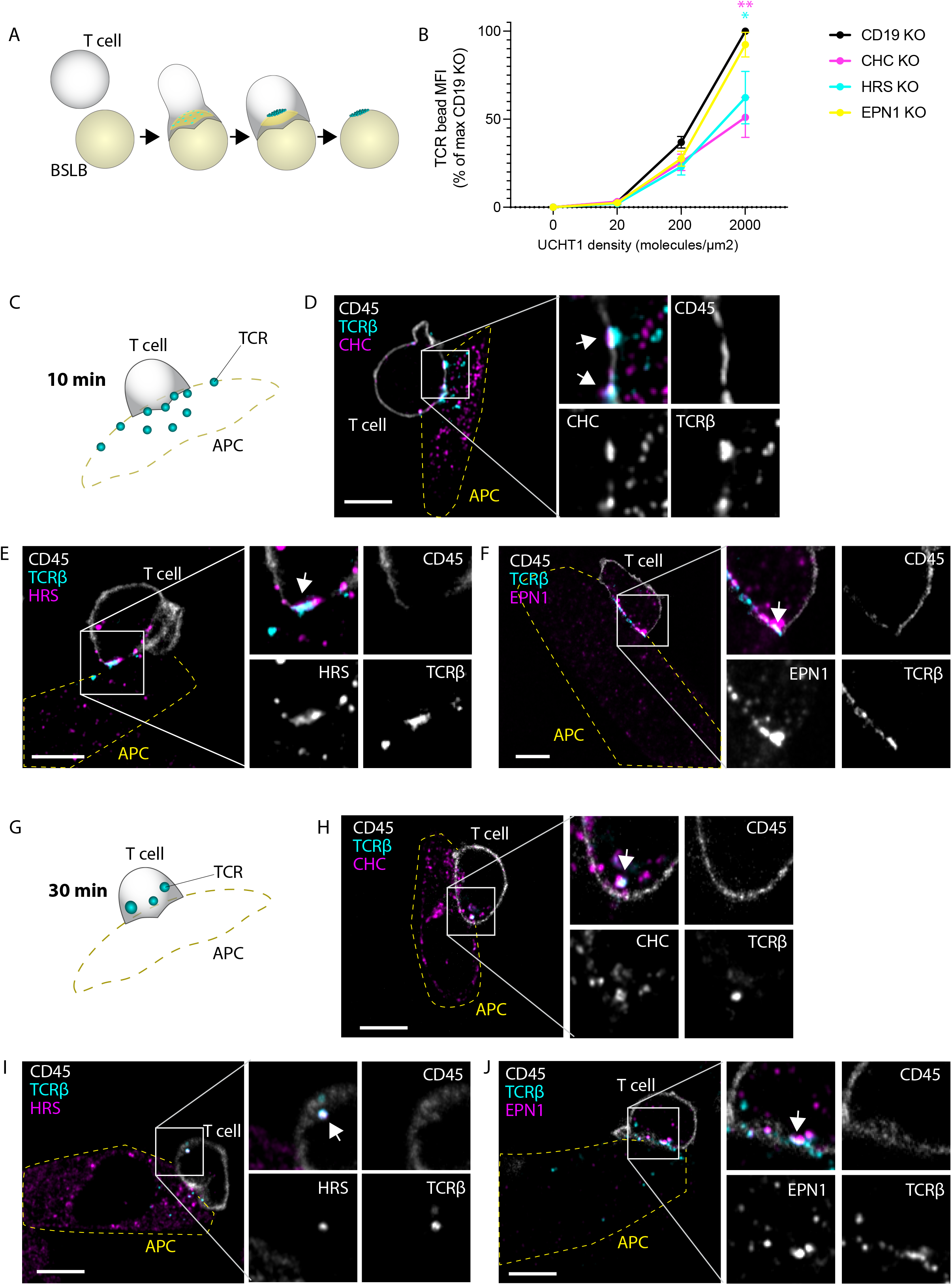
CHC and HRS are required for transfer of TCR loaded vesicles and they are recruited to TCR loaded endosomes. (**A**) Schematic of the vesicular transfer from T cells to Bead Supported Lipid Bilayers. ICAM-1 is labelled yellow and TCR is labelled cyan. As the TCR microclusters reach the center of the contact area, they are released as ectosomes which are left behind as the T cell detaches from the bead. (**B**) Quantification of the fluorescence intensity of beads with immunolabelled TCR transferred from hCD4 T cells at UCHT1 densities of 0, 20, 200 and 2000 molecules/μm^2^. The MFI represents the mean +/− SEM of 4 experiments relative to the max MFI of beads incubated with CD19 KO cells. *, p-value of < 0.05; **, p value of < 0.01 with unpaired twotailed non-parametric Mann Whitney test. (**C-J**) Airyscan micrographs of mCD4 AND T cells incubated with CHO-I-E^k^ antigen presenting cells for 10 min (**C-F**) and 30 min (**G-J**) and immunolabelled with anti-CD45, anti-TCRβ and either anti-CHC, anti-HRS or anti-EPN1. The outline of the APCs is indicated with a yellow dashed line. Scale bar, 5 μm.

To confirm that clathrin, HRS and EPN1 recruitment to the immunological synapse also occurs between T cells and antigen presenting cells, we incubated CHO cells expressing the I-E^k^-MCC peptide-MHC complex (CHO-I-E^k^-MCC) with AND mCD4 T cells for 10 and 30 minutes before we fixed and permeabilized them. We then immunolabelled the cells with antibodies targeting each of the proteins, together with H57-Fab to label TCRβ and anti-CD45 to label the T cells. Airyscan microscopy of the labelled cells revealed that clathrin, HRS and EPN1 were all recruited to the synaptic interphase and colocalized with TCRβ after 10 minutes (Figure 4C-F). Note that the T cells had already released TCR onto the APCs at this timepoint and that CD45 is excluded from TCR release sites.

After 30 min, most of the TCR was either transferred to the APCs or internalized by the T cell (Figure 4G-J). This was accompanied by a remarkable shift in localization of HRS and clathrin on the T cell side, a large fraction of which now associated with the TCR loaded internalized vesicles likely representing early/late endosomes. EPN1 on the other hand still appeared to associate with TCR at the plasma membrane.

When we also directly immunolabelled I-E^k^-MCC, we could observe that the internalized TCR colocalized extensively with the pMHC complex, thus indicating that the endosomal structures represent cointernalized antigen-TCR conjugates (Figure S4A). Some of these structures appeared to originate from microvillar protrusions from the APC interacting with the T cell body distal from the T cell-APC interphase which colocalized with CHC staining on the TCR side (Figure S4B). Similar protrusions originating from the T cell also reached distant APCs and other T cells, as previously reported by Kim *et al* ^7^. As shown in Figure S4C-D, HRS and CHC could be found at the base of such structures, thus suggesting that these proteins might also regulate this form of membrane transfer. Figure S4C shows how an individual AND T cell can form 4 different types of interactions with APCs and other T cells simultaneously, all of which are marked by HRS on the T cell side: 1. Primary synapse, 2. Secondary synapse, 3. Microvillar protrusion from the APC, 4. Microvillar protrusion from the T cell.

We then did CRISPR/Cas9-mediated KO of CD19, CHC, HRS and EPN1 in the AND cells and incubated them with CHO-I-E^k^ cells for 30 minutes before we fixed and permeabilized them as before. We immunolabelled TCRβ, I-E^k^-MCC and CD45 and used Imaris software to create a 3D mask of the T cells based on the CD45 signal (Figure 5A). This enabled us to segment I-E^k^-MCC positive vesicles internalized by the T cells (Figure 5B-C). When we counted the number of such vesicles per T cell, we could observe a median decrease of 50% following KO of CHC, HRS and EPN1 (Figure 5D). When we then quantified the sum intensity of I-E^k^-MCC per vesicle, we could observe a significant reduction following EPN1 KO (Figure 5E). The drop in intensity was apparently caused by a reduction in the volume of the internalized vesicles, as this parameter was also affected by EPN1 KO, but not by depletion of CHC or HRS (Figure 5F). We then segmented extracellular vesicles released by the T cells based on their H57 signal, disregarding any signal originating from within the T cell body (Figure 5G). When we counted the number of released vesicles per T cell, we could observe a strong reduction following CHC and HRS KO, but not after EPN1 depletion (Figure 5H).

**Figure 5.**
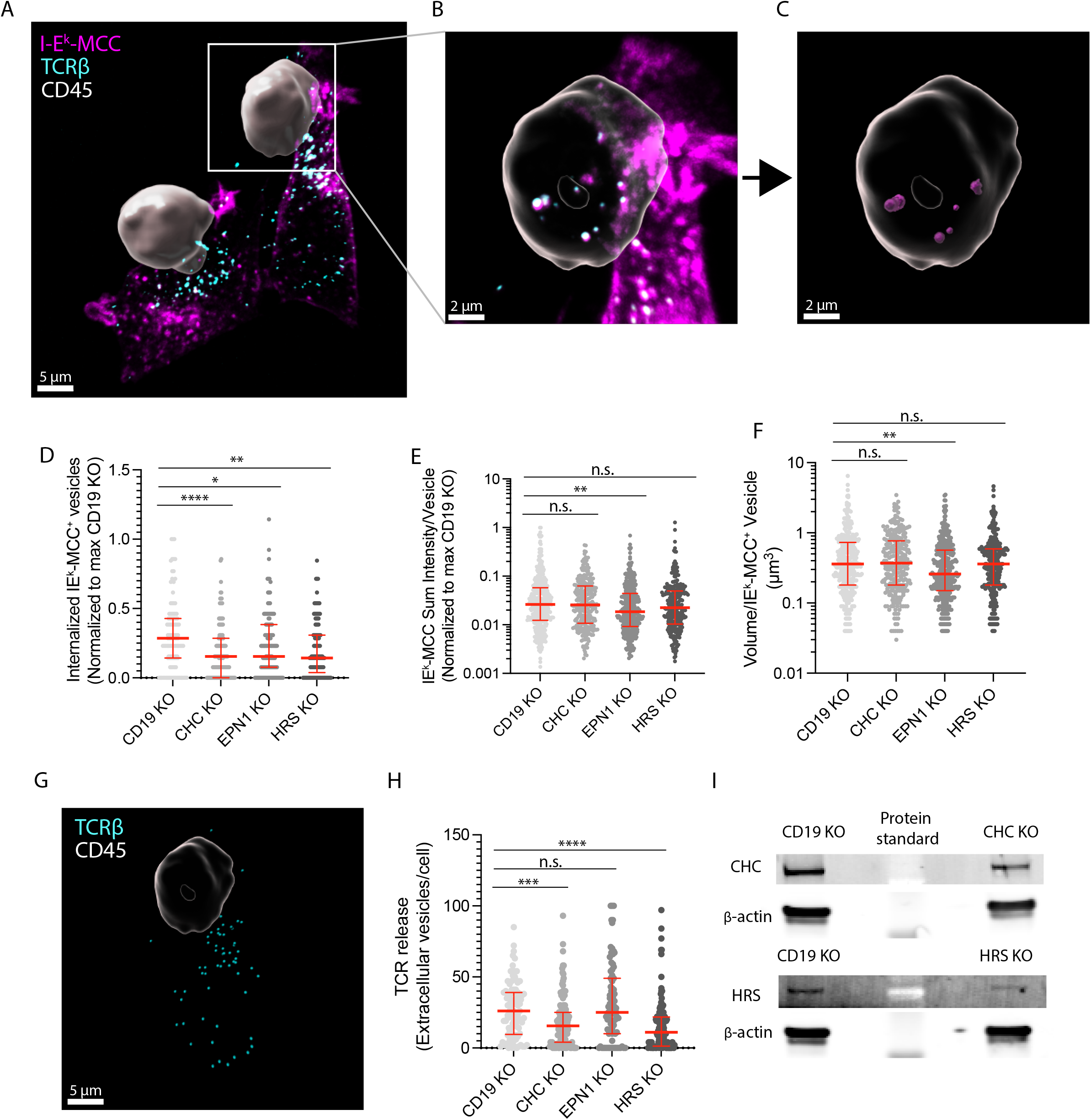
Clathrin, HRS and EPN1 control pMHC-TCR uptake. (**A-C**) Imaris 3D reconstructed micrographs of spinning disc confocal Z-stacks with a step size of 250 nm of mCD4 AND T cells incubated with CHO-I-E^k^ APCs for 30 min and immunolabelled with anti-CD45 (white), anti-TCRβ (cyan) and anti-I-E^k^-MCC (magenta). The CD45 signal has been used to create a mask of the T cell. (**D-F**) Quantification of the number of internalized I-E^k^-MCC positive vesicles per cell (**D**), sum I-E^k^-MCC fluorescence intensity per vesicle (**E**) and volume per I-E^k^-MCC positive vesicle (**F**) following knockout of CD19, CHC, EPN1 or HRS in mCD4 AND cells incubated for 30 min on CHO-I-E^k^ APCs. *, p-value of < 0.05; **, p value of < 0.01; ****, p value of < 0.0001 with unpaired two-tailed non-parametric Mann Whitney test. N_cells_ ≥ 111 per condition. Lines represent median value and error bars represent interquartile range. (**G**) Segmented TCR positive vesicles (cyan) released from one of the mCD4 T cells from (**A**). (**H**) Quantification of the number of released TCR positive vesicles per cell of the mCD4 AND T cells from (**D**). ***, p value of < 0.001; ****, p value of < 0.0001 with unpaired two-tailed non-parametric Mann Whitney test. Lines represent median value and error bars represent interquartile range. (**I**) A representative western blot of the protein levels of CHC, HRS and β-actin from one of the experiments in (**D**) following CHC and HRS KO, respectively.

Taken together, these data show that CHC, EPN1 and HRS are all involved in pMHC-TCR internalization on the T cell side, while only CHC and HRS are involved in release of TCR positive extracellular vesicles.

## Discussion

The role of clathrin in CD4^+^ T cell activation has thus far been studied and interpreted in a paradigm in which the secretory nature of the immunological synapse was yet to be discovered. Hence, even though several studies have identified clathrin as being involved in T cell activation, its functional relevance has not been fully understood. Here we show that clathrin is essential for release of TCR-loaded vesicles into the synaptic cleft and uptake of antigen-ligated TCR from the synapse into the T cell.

We show that clathrin is recruited to TCR microclusters by HRS and STAM2 where it mediates release of TCR loaded synaptic ectosomes into the synaptic cleft. This initiates a temporal transition during which the endosomal clathrin adaptor EPN1 initiates clathrin-mediated internalization of antigen ligated to TCR. Depletion of EPN1, HRS or clathrin blocks pMHC-TCR endocytosis. AP2 is present at the synaptic interphase but is not specifically recruited to TCR microclusters and seems redundant for TCR release or retrieval.

The decrease in antigen uptake following HRS KO might be caused by the block in TCR release and CHC arrest at the cSMAC as we show after HRS and TSG101 KO in human CD4 T cells activated on supported lipid bilayers. However, we cannot exclude the possibility of HRS being directly involved in pMHC-TCR endocytosis.

Several studies have indicated that clathrin is not involved in uptake of activated TCR ^20–22^. However, those studies have relied on indirect analysis of CD3 expression on the cell surface or on quantification of internalized TCR positive vesicles near the synapse and not direct analysis of total antigen positive vesicles. We show that antigen can be internalized far from the T cell-APC interphase. Furthermore, some of those studies have been conducted in systems where T cells have been activated on inert substrates (i.e. antigen coated plastic surfaces), from which the T cells might not be able to internalize antigen-TCR conjugates. Indeed, we also fail to detect vast uptake of TCR conjugated to UCHT1 from SLBs. However, we observe clear defects in pMHC-TCR uptake after CHC KO when directly counting the number of antigen positive vesicles internalized by T cells following activation on APCs.

EPN1 has been reported to associate with polyubiquitinated receptors but has low affinity for monoubiquitin ^31^. HRS and STAM2 on the other hand have been shown to associate stably with monoubiquitinated cargos ^32^. Activated TCRs transitioning from mono-to poly-ubiquitination during synapse maturation might thus determine whether the receptors are released in synaptic ectosomes or internalized and degraded. Polyubiquitination of CD3ζ has been shown to be required for TCR signaling termination following activation by anti-CD3ε, although this appeared to be independent of global receptor downregulation ^33^. Whether internalization of TCR directly conjugated to pMHC on APCs requires polyubiquitination is not known. This could however explain the time delay in EPN1 recruitment compared to HRS and STAM2 to TCR microclusters as well as our observation of rapid release of TCR onto antigen presenting cells and delayed internalization of antigen-conjugated TCR.

We conclude that TCR loaded vesicles are rapidly released directly from the plasma membrane upon T cell activation in a process mediated by clathrin and the ESCRT machinery. There is no specific term for the process, described here, of clathrin and ESCRT dependent evagination of plasma membrane into small extracellular vesicles. Formation of large vesicles at the plasma mebrane can take place through a process of blebbing ^34^. Given the precedent of refering to small vesicles that bud form the plasma membrane as ectosomes ^11^, we suggest *ectocytosis* as a term to describe this process. Following a temporal shift in clathrin adaptor recruitment, EPN1 and clathrin mediated internalization of antigen-ligated TCR is initiated. Although this clathrin dependent internalization dominates retrieval of cSMAC associated TCR in our system, we cannot exclude the possibility of clathrin-independent uptake of triggered TCR working in parallel with the clathrin pathway. It is also possible that the antigen potency, co-stimulatory signals, and/or antigen presenting cell mechanics dictate the internalization mechanism triggered.

Nevertheless, the data presented in this manuscript establishes clathrin and the ESCRT machinery as essential players in regulating direct release and retrieval of triggered TCR at the immunological synapse.

## Materials and Methods

### Cell culture

#### Human T cells

Peripheral blood from healthy donors was acquired from the National Health Service blood service under ethics license REC 11/H0711/7 (University of Oxford). CD4^+^ cells were isolated by negative selection (RosetteSepTM Human CD4^+^ T cell Enrichment Cocktail, STEMCELL technologies, Cambridge, UK; #15023) following the manufacturer’s protocol. Donor specific CD4^+^ T cells were plated at 1×10^6^ cells/ml with CD3/CD28 activation beads (Dynabeads, ThermoFisher Scientific, Loughborough, UK; #11132D) at a 1:1 ratio with 50 U/ml IL-2 at 2 ml per well in a 24 well plate and expanded for three days at 37°C in a CO_2_ incubator. On day three, cells were resuspended, counted and replenished with fresh media to reach the original density of 1×10^6^ cells/ml. The cells were expanded for four more days replenished with 50 U/ml IL-2 and fresh culture media every second day. The cells were left without IL-2 the day before they were incubated on SLBs.

#### Murine T cells

AND Mouse T cells were generated from spleen and lymph nodes of TcrAND B10.Br mice; mice were housed under pathogen-free conditions in the Kennedy Institute of Rheumatology Animal Facility in accordance with local and Home Office regulations. The mice were housed in individually ventilated cages (IVC) with corn cob bedding. 12 hours of light/dark cycle with half an hour of dim light period in place. Appropriate environmental enrichment was provided: Enviro-dri, and housing tunnel, cage balcony and chew blocks. The temperature was maintained at 21 degrees +/− 2 and the humidity 55% +/− 10. The protocols were reviewed by local Veterinary Surgeon (Vet) and Named Animal Care and Welfare Officer (NACWO) before being reviewed and approved by Animal Welfare and Ethical Review Body (AWERB). All procedures were conducted in accordance with the UK Scientific Procedures Act of 1986 and overlooked by University of Oxford Department of Biomedical Services, Clinical medicine ORC (old road campus).

AND T cells were plated at 1×10^7^ cells/ml with 1 μM MCC peptide with 50 U/ml IL-2 at 2 ml per well in a 24 well plate and expanded for three days at 37°C in a CO_2_ incubator. On day three, cells were resuspended, counted and replenished with fresh media to reach a density of 1×10^6^ cells/ml. The cells were expanded for four more days replenished with 50 U/ml IL-2 and fresh culture media every second day. The cells were left without IL-2 the day before they were used in downstream procedures (AND T cell blasts). Mouse TCRβ was labelled with the Fab fragment of anti-Mouse TCRβ clone H57-597 (Biolegend; #109201).

Chinese hamster ovary (CHO) cells stably expressing I-E^k^-MCC (CHO-I-E^k^) were maintained by passage every 3 days in complete RPMI 1640 medium (ThermoFisher Scientific, Loughborough, UK; #31870074).

### Planar Supported Lipid Bilayer Experiments

Flow chambers (sticky-Slide VI 0.4, Ibidi, Thistle Scientific LTD, Glasgow, UK; #80608) were attached to cleanroom cleaned coverslips (SCHOTT UK Ldt, Stafford, UK; #1472315) and cured for thirty minutes. 50ul of 12.5 mol% DOGS-NTA(Ni) lipids in DOPC was loaded in each IBIDI channel and incubated for 20 minutes. All lipids were obtained from Avanti Polar Lipids (Alabaster, AL). The channels were washed three times with 200ul of 0.1% BSA/HBS (20 mM HEPES, 137 mM NaCl, 1.7 mM KCl, 0.7 mM Na_2_PO_4_, 5 mM glucose, 2 mM MgCl_2_, 1 mM CaCl_2_, pH 7.2) and blocked with 200ul of 2% BSA/HBS for 20 minutes. The channels were washed again three times with 200ul of 0.1% BSA/HBS before 200ul of protein mix (ICAM1-AF405, UCHT1-AF488 or I-E^k^-MCC) in 0.1% BSA/HBS was loaded and incubated for 20 minutes. The protein concentrations were calibrated by flow cytometry to yield the desired density (described below). The channels was washed again three times with 200ul of 0.1% BSA/HBS. Fluorescence Recovery After Photobleaching (FRAP) was then performed to control bilayer mobility on an inverted Olympus FV1200 confocal microscope. All incubation steps were done at room temperature.

1 x 10^4^ T cells were addedd to each channel and incubated at 37°C for 5 or 20 minutes. Cells were fixed with 100ul of 4% Paraformaldehyde (PFA) in PHEM buffer (60mM PIPES, 25mM HEPES, 10mM EGTA, and 4mM MgSO4·7H20) for 10 minutes, washed and permeabilised with 100ul of 0.1% Triton X-100 in 0.1% BSA/HBS for 2 minutes. The channels were washed three times with 200ul PBS before blocking solution with 5% goat serum or BSA in PBS was added for one hour. Antibodies was then diluted in 200 μl blocking solution and incubated with the cells overnight at 4°C. Each channel was then washed 3 times with 200 μl PBS before the appropriate secondary antibody was added and incubated for 1 hr. The channels were washed again 3 times with 200 μl PBS.

TIRFM imaging was performed either with an Olympus IX83 inverted microscope (Keymed, Southend-on-Sea, UK) equipped with a 150x 1.45 NA oil immersion objective and an EMCCD camera (Evolve Delta, Photometrics, Tucson, AZ) or a Deltavision OMX V4 system (Applied Precision, GE Healthcare) equipped with a 60x ApoN NA 1.49 objective (Olympus), and three cooled sCMOS cameras (PCO). For live-cell imaging, the stage and objective were temperature controlled at 37°C and the sample holder was enclosed by a humidified 5% C02 incubation chamber.

### CRISPR/Cas9-mediated KO

Freshly isolated CD4^+^ T cells were activated for 3 days on CD3/CD28 activation beads (Dynabeads ThermoFisher Scientific, Loughborough, UK; #11132D) as before. They were then washed three times in Opti-MEM (Gibco; #11058021). 1ul 200 uM cripsrRNA was mixed with 1ul 200 uM tracrRNA in an RNAse-free PCR tube and incubated at a Thermal Cycler C1000 (Bio Rad) at 95°C for 5 min and allowed to cool to RT slowly. 7.5 μl of 20μM Cas9 was added to the guideRNA complex at room temperature while swirling the pipette to mix the reagents. The mix was then incubated at the Thermal Cycler at 37°C for 15 min and slowly cooled to room temperature. Finally 1ul of 200uM electroporation enhancer was added to the RNP mix at RT and gently mixed. All reagents were from IDT DNA.

### T cell activation on CHO-I-E^k^ cells

1×10^4^ CHO-I-E^k^ cells were plated in each channel of 6 channel flow chambers (μ-Slide VI 0.4, Ibidi, Thistle Scientific LTD, Glasgow, UK; # 80606) for 24h in complete RPMI 1640 medium (ThermoFisher Scientific, Loughborough, UK; #31870074) before 1×10^4^ AND T cell blasts were added for 10 or 30 minutes. These were subsequently fixed with 100ul of 4% PFA in PHEM buffer (60mM PIPES, 25mM HEPES, 10mM EGTA, and 4mM MgSO4·7H20) for 10 minutes, washed and permeabilised with 100ul of 0.1% Triton X-100 in PBS for 2 minutes. The channels were washed three times with 200ul PBS before blocking solution with 5% BSA in PBS was added for one hour. Unconjugated primary antibodies were then diluted in 200 μl blocking solution and incubated with the cells overnight at 4°C. Each channel was then washed 3 times with 200 μl PBS before the appropriate secondary antibody was added and incubated for 1 hr. The channels were washed again 3 times with 200 μl PBS before Z-stacks were aquired either with a Zeiss AiryScan 880 confocal microscope or a Nikon Ti2-E microscope with a spinning disk.

### Bead Supported Lipid Bilayers (BSLB)

1×10^6^ non-functionalized silica beads (5.0 +/− 0.05 μm diameter, Bangs Laboratories, Inc., #) per condition were washed three times with PBS in 1.5 ml conical microcentrifuge tubes. Then, BSLBs were formed by incubating washed silica beads with 12.5 mol% DOGS-NTA(Ni) DOPC. All liposome stocks mixed for the formation of SLB were used at a final lipid concentration of 0.4 mM. Then, to remove excess lipids, the freshly formed BSLB were washed three times with 1% HSA/HBS. BLSB were then blocked with 5% BSA, 100 μM NiSO4 in PBS for no longer than 20 minutes.

Protein dilutions for planar SLB and downstream BSLB procedures were calibrated by making serial 2x dilutions of the desired protein from 1/100 to 1/50200. 100μl of each dilution was added to the beads and incubated for 20 minutes on a shaker at 800 rpm before the beads were washed three times in 1000ul of 0.1% BSA/HBS and analyzed by flow cytometry. Standard calibration curves were calculated using Quantum Alexa Fluor 647 MESF (Bangs Laboratories, Fishers, IN; # 647-A) or Quantum Alexa Fluor 488 MESF (Bangs Laboratories, Fishers, IN; #488-A).

Prior to incubation with BSLBs, T cells were washed two times with fully supplemented Phenol Red-free RPMI 1640 lacking IL-2 and resuspended to an assay concentration of 2.5 x 10^6^ cells/mL. Then, T cells (2.5 x 10^5^/well) were incubated with BSLB at 1:1 ratio in U-bottom 96 well plates (Corning, #) for 90 min at 37°C and 5% CO_2_. For gentle dissociation of BSLB:cell conjugates, culture plates were gradually cooled down by incubation at RT for 15 min, followed by incubation on ice for a minimum of 40 min. Then, cells and BSLB were centrifuged at 500 x g for 5 min and then gently resuspended in ice-cold 5% BSA in PBS prior to staining for flow cytometry analysis.

### Measurement of Synaptic Transfer by conventional Flow Cytometry

Staining with fluorescent dye conjugated antibodies was performed immediately after dissociation of cells and BSLB conjugates. Staining was performed in ice-cold 5% BSA in PBS pH 7.4 (0.22 μm-filtered) for a minimum of 30 min at 4°C and agitation to avoid BSLB sedimentation (700 rpm in the dark). Then, cells and BSLB were washed three times and acquired immediately using an LSR Fortessa X-20 flow cytometer equipped with a High Throughput Sampler (HTS). For absolute quantification, we MESF beads which were first acquired to set photomultiplier voltages to position all the calibration peaks within an optimal arbitrary fluorescence units’ dynamic range (between 10^1^ and 2 x 10^5^, and before compensation). Fluorescence spectral overlap compensation was then performed using single colour-labelled cells and BSLB, and unlabelled BSLB and cells. For markers displaying low surface expression levels unstained and single colour stained UltraComp eBeads (Thermo Fisher Scientific Inc.; #01-2222-42) were used for the calculation of compensation matrixes. After application of the resulting compensation matrix, experimental specimens and Quantum MESF beads were acquired using the same instrument settings. In most experiments acquisition was set up such as a minimum of 2 x 10^4^ individual BSLB were recorded. To reduce the time of acquisition of high throughput experiments a minimum of 1 x 10^4^ single BSLB was acquired per condition instead.

## Data Analysis

Automated segmentation of individual cells from each TIRF micrograph was performed with FIJI (version 2.3.0/1.53f) based on the brightfield channel, followed by background subtraction (rolling ball, 50 pixel radius). Calculation of relative fluorescence intensity was done using using Matlab (version 2021b). 3D segmentation and quantification of confocal Z-stacks was done using Imaris batch analysis. For radial averaging, the Fiji macro https://github.com/donFellus/radAv was used. This macro is specifically designed to analyze the average position of subcellular structures within the immunological synapse. It rotates a selected image 360 times 1 degree and creates a copy following each rotation. It then merges all the copies to create a radial positional average of the fluorescence signals in the original image.

Statistical analysis and plots were generated using GraphPad Prism (version 9.2.0). Analysis of data generated by Flow cytometry was performed with FlowJo LLC (version 10.8).

## Acknowledgments

This work was supported by The Research Council of Norway in conjunction with Marie Sklodowska-Curie Actions 275466, the Kennedy Trust for Rheumatology Research, the Wellcome Trust (100262Z/12/Z) and the European Commission (ERC-2014-AdG_670930).

---

**Figure S1.**
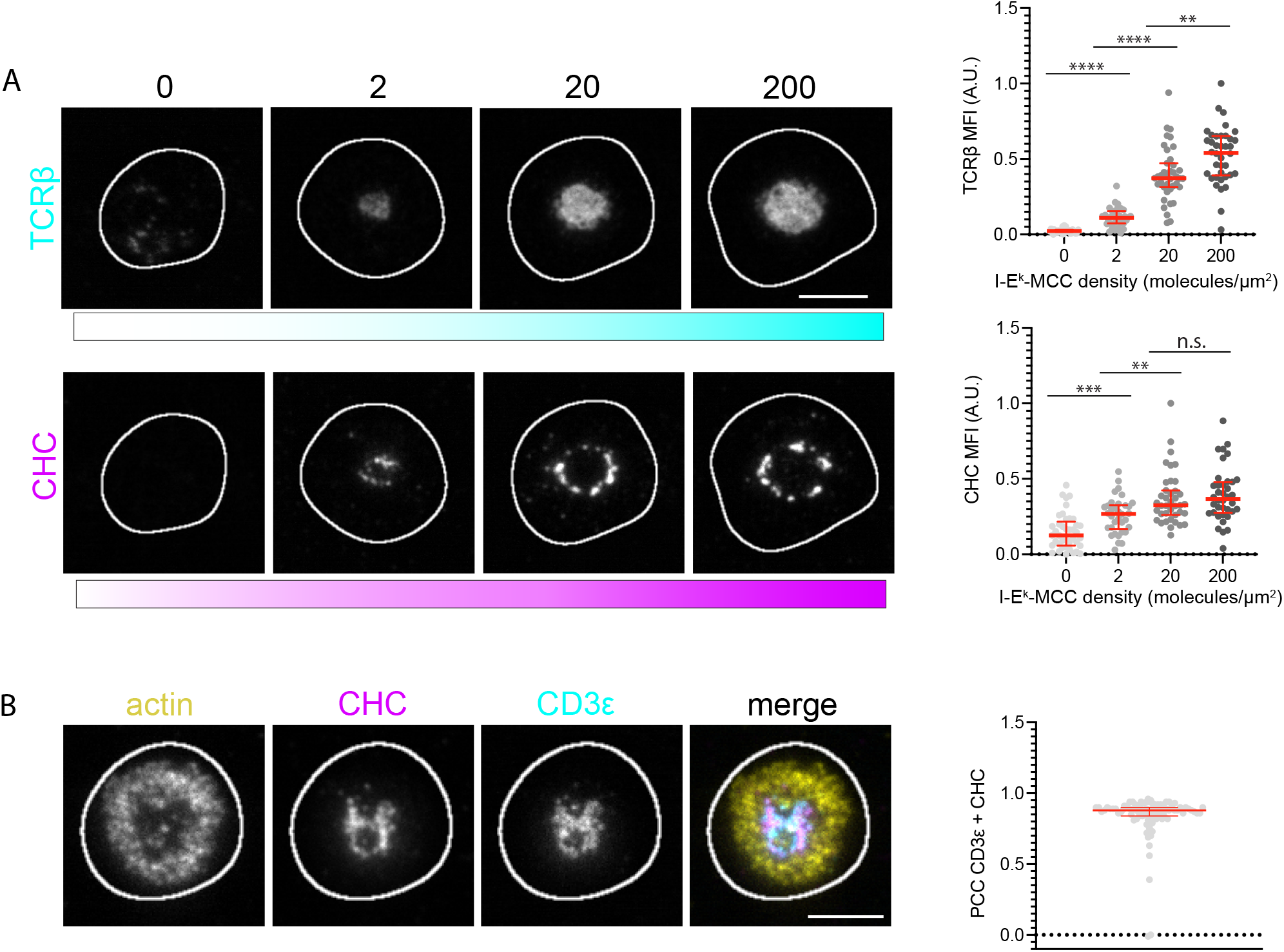
Clathrin is recruited to the immunological synapse, related to Figure 1. (**A**) Representative TIRF micrographs of AND mCD4 T cells incubated on SLBs either with ICAM-1-AF405 (200/μm^2^) alone for 5 min or with ICAM-1-AF405 + I-E^k^-MCC (2, 20 or 200/μm^2^) for 20 min, labelled with anti-mouse TCRβ (cyan) and anti-CHC (magenta). N_cells_ > 34 per density. Scale bar, 5 μm. Right panels are quantifications of the MFI of TCRβ and CHC across the synaptic interphase. **, p value < 0.01; ***, p value < 0.001; ****, p value of < 0.0001 with unpaired two-tailed non-parametric Mann Whitney test. Lines represent median value and error bars represent interquartile range. (**B**) Representative TIRF micrographs of a hCD4 T cell incubated on a SLB with ICAM-1-AF405 (200/μm^2^) + UCHT1-AF488 (engages CD3ε, cyan, 30/μm^2^) for 20 min, labelled with anti-CHC (magenta) and phalloidin-AF647 (yellow). The right panel is quantification of the PCC between CHC and CD3ε across the synaptic interphase. N_cells_ = 97. Scale bar, 5 μm. Lines represent median value and error bars represent interquartile range.

**Figure S2.**
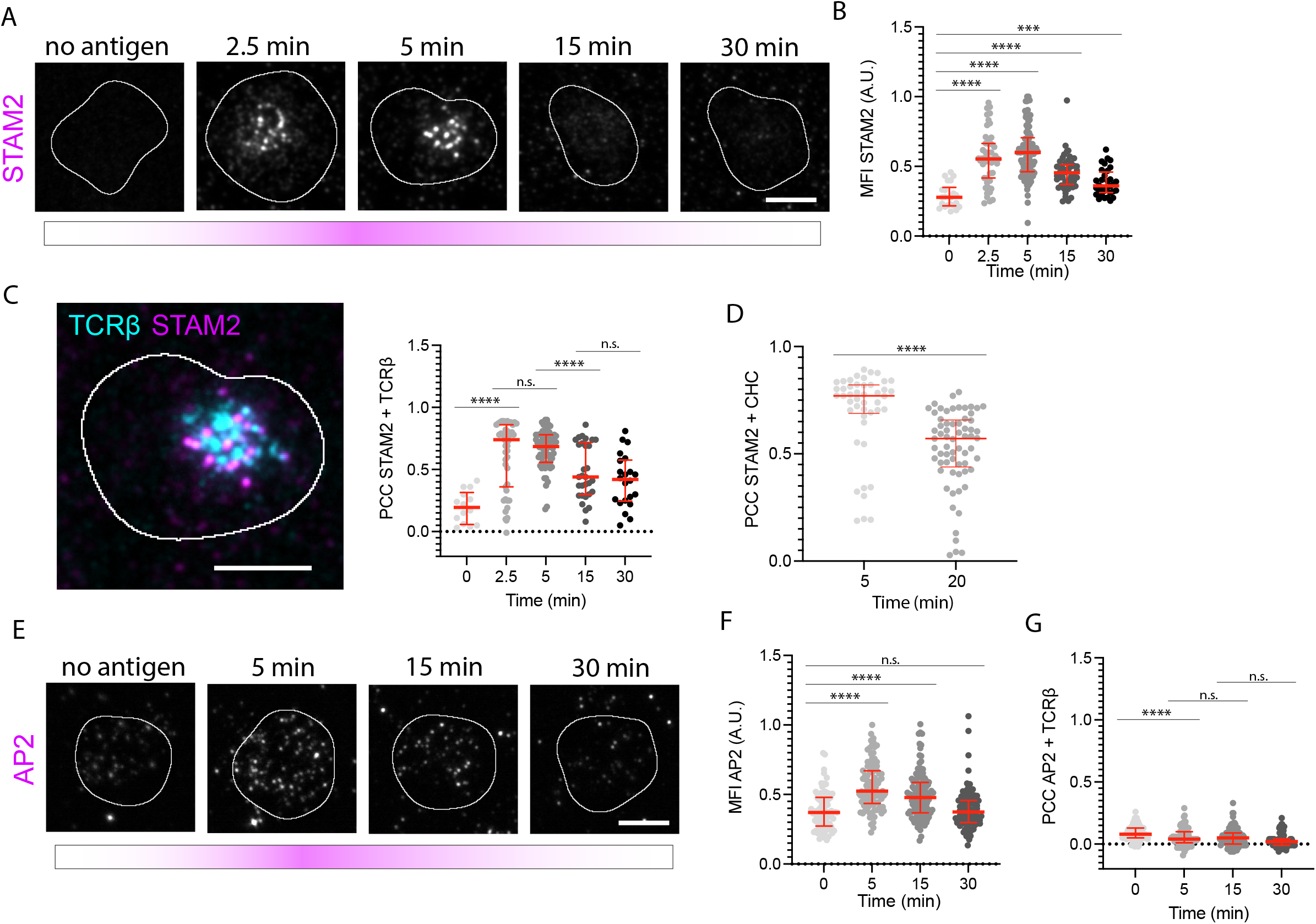
STAM2 and AP2 are recruited to the immunological synapse, but AP2 is not recruited to TCR microclusters, related to figure 2. (**A**) Representative TIRF micrographs of AND mCD4 T cells incubated on SLBs either with ICAM-1-AF405 (200/μm^2^) alone for 5 min or with ICAM-1-AF405 + I-E^k^-MCC (20/μm^2^) for 2.5, 5, 15 and 30 min, labelled with anti-STAM2 (magenta). N_cells_ ≥ 48 per timepoint. Scale bar, 5 μm. (**B**) Quantification of the temporal MFI of STAM2 across the synaptic interphase from the micrographs in (**A**). ***, p value < 0.001; ****, p value of < 0.0001 with unpaired two-tailed non-parametric Mann Whitney test. Lines represent median value and error bars represent interquartile range. (**C**) Representative TIRF micrograph of an AND mCD4 T cell incubated on SLBs either with ICAM-1-AF405 (200/μm^2^) + I-E^k^-MCC (20/μm^2^) for 5 min, labelled with anti-STAM2 (magenta) and anti-TCRβ (cyan). The right panel is quantification of the temporal PCC between STAM2 and TCRβ across the synaptic interphase. Scale bar, 5 μm. ****, p value of < 0.0001 with unpaired two-tailed non-parametric Mann Whitney test. Lines represent median value and error bars represent interquartile range. (**D**) Quantification of the temporal PCC between STAM2 and CHC. N ≥ 46 per timepoint. ****, p value of < 0.0001 with unpaired two-tailed non-parametric Mann Whitney test. Lines represent median value and error bars represent interquartile range. (**E**) Representative TIRF micrographs of AND mCD4 T cells incubated on SLBs either with ICAM-1-AF405 (200/μm^2^) alone for 5 min or with ICAM-1-AF405 + I-E^k^-MCC (20/μm^2^) for 5, 15 and 30 min, labelled with anti-mouse TCRβ (cyan) and anti-AP2 (magenta). N_cells_ ≥ 87 per timepoint. Scale bar, 5 μm. (**F-G**) Quantification of the temporal MFI of AP2 (**F**) and the temporal PCC between AP2 and TCRβ (**G**) across the synaptic interphase from the micrographs in (**E**). ****, p value of < 0.0001 with unpaired two-tailed non-parametric Mann Whitney test. Lines represent median value and error bars represent interquartile range.

**Figure S3.**
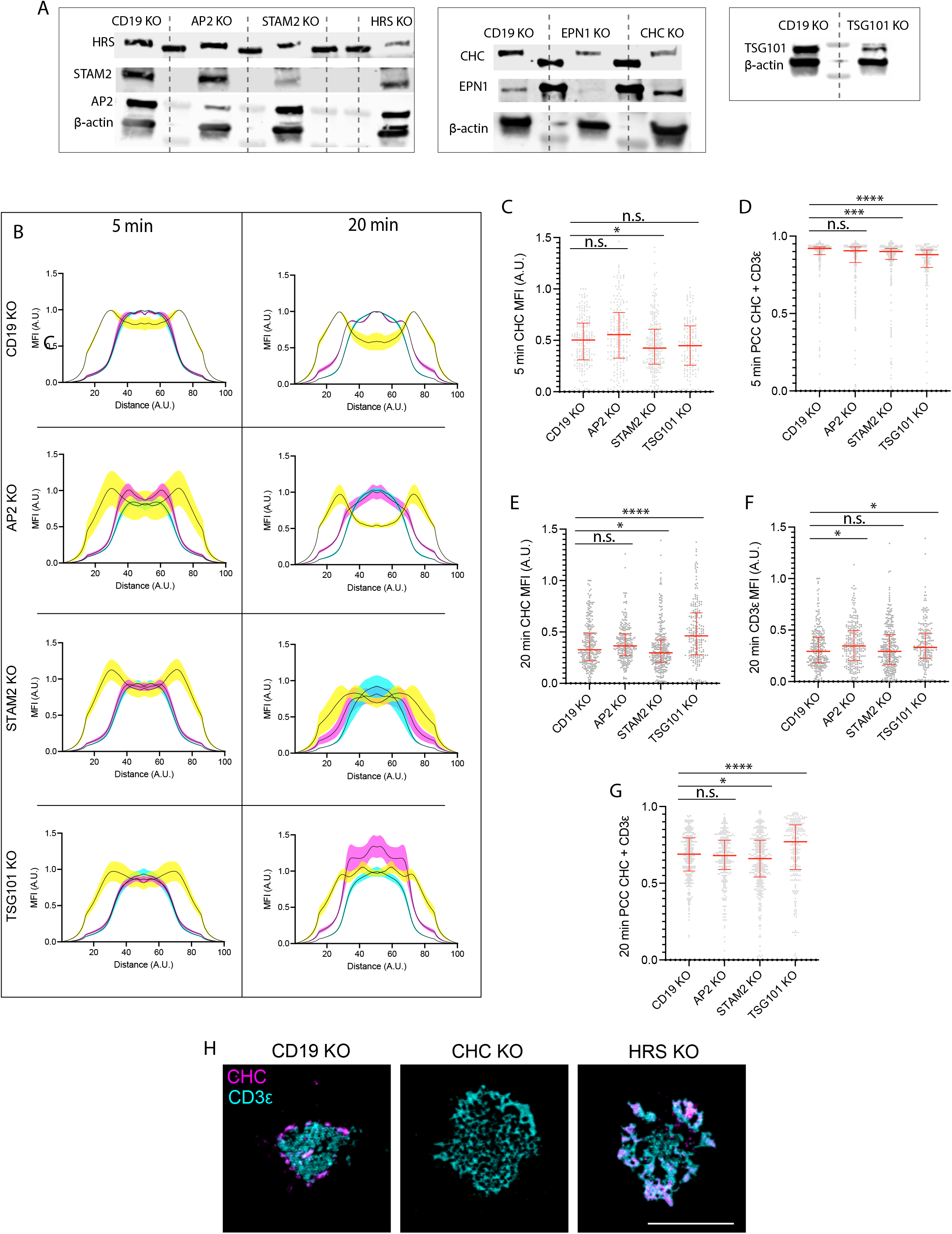
CHC, HRS, STAM2 and TSG101 are required for immunological synapse formation, related to figure 3. (**A**) Representative western blots of the protein levels of CHC, HRS, EPN1, STAM2, AP2 and TSG101 following CRISPR/Cas9-mediated KO of the respective proteins. Anti-β-actin was included as loading control. Dashed lines indicate protein standard. (**B**) Radial averages of hCD4 CD19, AP2, STAM2 and TSG101 KO T cells incubated on the bilayers from (**Figure 3B**) for 5 minutes (**A**) and 20 minutes (**B**). MFI represents mean fluorescence intensity from 3 experiments +/− SEM. (**C-G**) Quantification of the MFI of CHC (**C, E**), CD3ε (**F**) and the PCC between CHC and CD3ε (**D, G**) across the synaptic interphase from hCD4 CD19, CHC, HRS and EPN1 KO T cells incubated on the bilayers from (**B**) for 5 and 20 minutes, respectively. *, p-value of < 0.05; ***, p value of < 0.001; ****, p value of < 0.0001 with unpaired two-tailed non-parametric Mann Whitney test. N_cells_ ≥ 165 per condition. Lines represent median value and error bars represent interquartile range. (H) Extended TIRF-SIM micrographs of hCD4 CD19, CHC and HRS KO T cells incubated on bilayers from **Figure 3B**. Scale bar, 5 μm.

**Figure S4.**
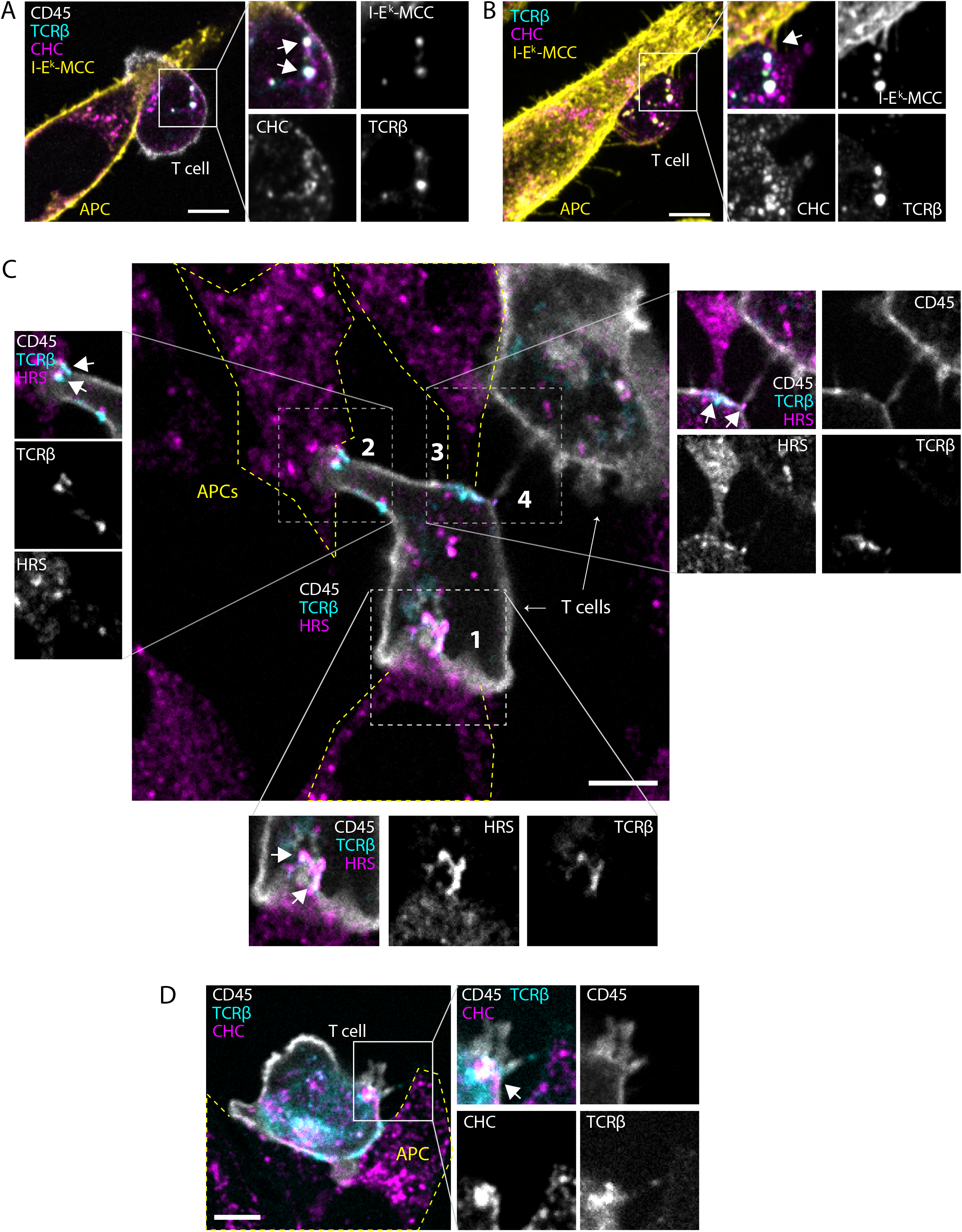
Clathrin and HRS regulate microvillar membrane transfer, related to figure 4. (**A-B**) A slice from an Airyscan^®^ Z-stack with a step size of 250 nm of an mCD4 AND T cell incubated with CHO-I-E^k^ antigen presenting cells for 30 min and immunolabelled with anti-CD45 (white), anti-TCRβ (cyan), anti-CHC (magenta) and anti-I-E^k^-MCC (yellow). **B** is a maximum intensity projection of the Z-stack from **A** emphasizing the microvillar protrusions from the APC. Scale bar, 5 μm. (**C-D**) A slice from a spinning disc confocal Z-stack with a step size of 250 nm of mCD4 AND T cells incubated with CHO-I-E^k^ antigen presenting cells for 5 min and immunolabelled with anti-CD45 (white), anti-TCRβ (cyan), anti-I-E^k^-MCC (yellow) and anti-HRS (**C**) or anti-CHC (**D**) (magenta). The intensities in some of the cropped images has been adjusted for emphasis. Yellow dashed lines indicate outlines of APCs. Scale bar, 5 μm.

